# The Architecture of GluD2 Ionotropic Delta Glutamate Receptor Elucidated by cryo-EM

**DOI:** 10.1101/2020.01.10.902072

**Authors:** Ananth Prasad Burada, Janesh Kumar

## Abstract

GluD2 receptors belong to the orphan delta receptor family of glutamate receptor ion channels. These receptors play key roles in synaptogenesis and synaptic plasticity and are associated with multiple neuronal disorders like schizophrenia, autism spectrum disorder, cerebellar ataxia, intellectual disability, paraplegia, retinal dystrophy, etc. Despite the importance of these receptors in CNS, insights into full-length GluD2 receptor structure is missing till-date. Here we report cryo-electron microscopy structure of the rat GluD2 receptor in the presence of calcium ions and the ligand 7-chlorokynurenic acid, elucidating its 3D architecture. The structure reveals a non-swapped architecture at the extracellular aminoterminal (ATD), and ligand-binding domain (LBD) interface similar to that observed in GluD1; however, the organization and arrangement of the ATD and LBD domains are unique. While our results demonstrate that non-swapped architecture is conserved in the delta receptor family, they also highlight the differences that exist between the two member receptors; GluD1 and GluD2.

## Introduction

GluD2 is a member of the orphan delta glutamate receptors and forms the fourth class of the ionotropic glutamate receptor (iGluR) family along with GluD1^1^. The other three classes being α-amino-3-hydroxy-5-methyl-4-isoxazole propionic acid (AMPA), kainate (KA) and N-methyl-D-aspartate (NMDA) receptors. Unlike AMPA, KA, and NMDA receptors ^2^ that are structurally very well characterized, structure-function relationships for delta receptors are still limited ^3^. GluD2 is expressed in multiple brain regions like the cerebral cortex, hippocampus, striatum, thalamus, mesencephalon, and retina at low levels ^4^. However, GluD2 expression levels are highest in cerebellar Purkinje cells (PCs), where it is known to form trans-synaptic complexes with presynaptic neurexins from the parallel fibers (PF) with the help of cerebelin1 (Cbln1)^5,6^. This ability of GluD2 to act as a synaptic organizer is vital for the maintenance of PF-PC synapses in the brain. Notably, GluD2 is associated with autism spectrum disorder, schizophrenia, cerebellar ataxia, intellectual disability, attention deficit hyperactivity disorder, and motor and cognitive deficit disorders ^7 8 9 10 11^.

While crystal structures of GluD2 amino-terminal domains (ATD) ^12^, ligand-binding domains (LBD) ^13^ and the intact extracellular region (ATD-LBD) ^14^ and ATD-Cbln1 ^12^ complex have been reported, the full-length structure of GluD2 is still lacking hindering further progress in the field.

In order to address this, we have determined the structure of homotetrameric rat GluD2 receptors in the presence of 1mM 7-Chlorokynurenic acid (7-CKA) and 1 mM Ca^2+^ ions using single-particle cryo-electron microscopy (cryo-EM). The structure reveals non-swapped architecture similar to GluD1 receptors and distinct from other iGluRs ^15^. However, notably, the ATD and LBD layer organization in GluD2 is unique and asymmetric. Our results not only provide insights into the architecture of orphan GluD2 receptors but also highlight structural differences within the delta receptor family.

## Results

We screened multiple C-terminally truncated rat GluD2 constructs via FSEC ^16^ and identified GluD2Δ870 for large-scale expression and purification from HEK293 GnTI^-^ cells in suspension using established protocols (**Fig. 1 A and B**). Purified receptor (GluD2Δ870) was complexed with 1mM 7-chlorokyneurenic acid (7-CKA) and 1mM Ca^2+^ ^17^ and subjected to cryo-EM analysis. Protein particles (~60000) were manually picked from micrographs and were cleaned up by iterative 2D classification (**Fig. 1 C**). *Ab-initio* 3D reconstruction followed by 3D refinement and other optical refinements as implemented in cryoSPARC v2 ^18^ (**Supplementary Fig. 1; Table 1**) for 3D refinement resulted into final map at ~7.9 Å resolution as estimated by the gold-standard FSC 0.143 criteria ^19^ (**Supplementary Fig. 2**). The 3D map had distinct features for the extracellular amino-terminal (ATD) and ligandbinding domain (LBD) that were modelled and refined (**Fig. 1 D**). However, the transmembrane domain was poorly resolved and hence was not fitted in the EM density map.

**Fig. 1:**
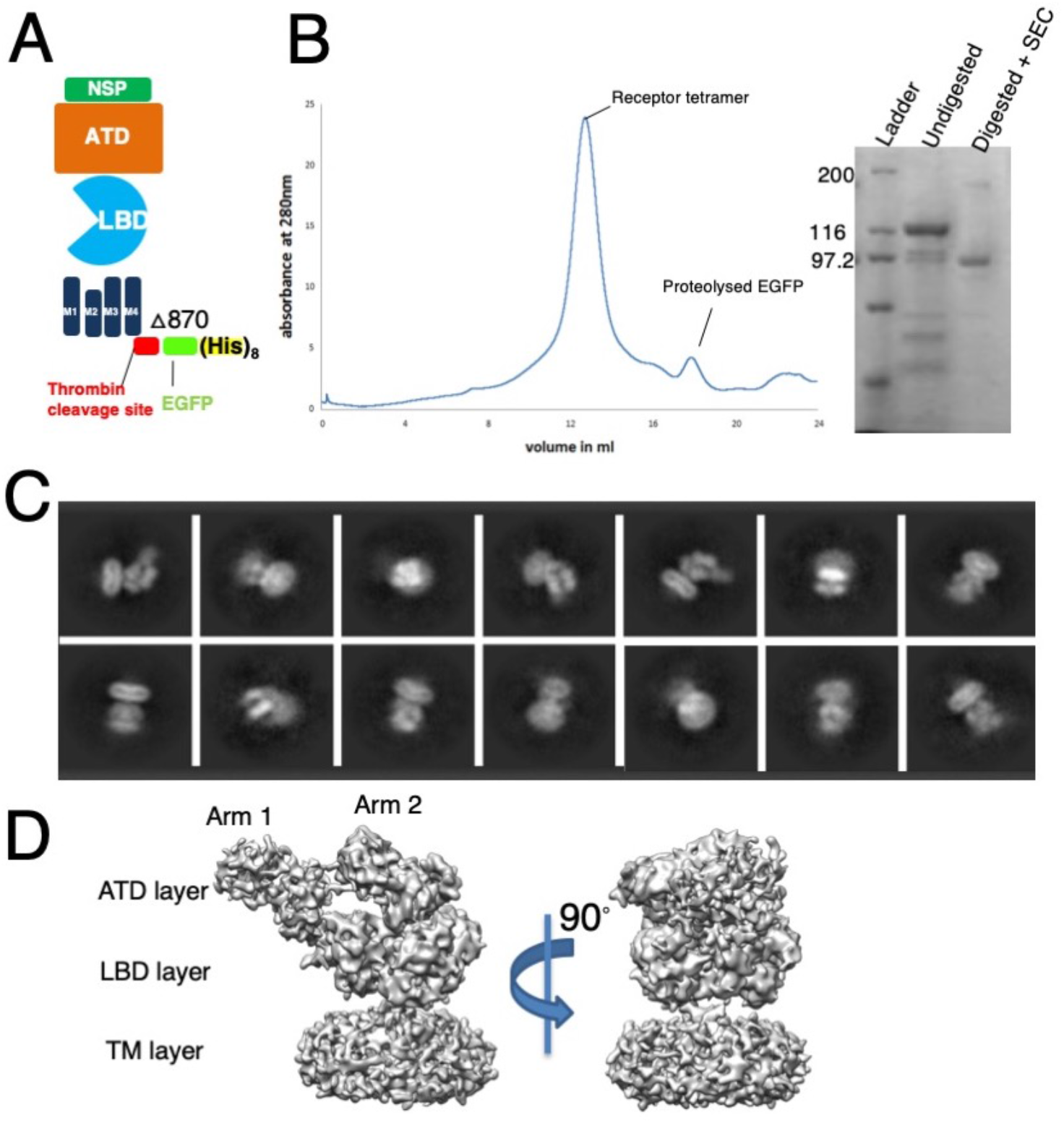
Construct design, purification, and Cryo-EM data processing for GluD2 receptors. **(A)** Schematic representation of the GLuD2 construct design. The C-terminal truncation at residue 870 and C-terminal thrombin cleavage site along with GFP and Octa histidine tag are indicated. **(B)** Size-exclusion profile of the purified GluD2 protein highlighting receptor stability in optimized buffer conditions. SDS-PAGE of the purified protein before and after thrombin digestion and SEC purification is shown in the inset. (**C)** Representative 2D class averages from reference-free 2D classification are shown. **(D)** The final refined 3D map of GluD2 is shown. Different receptor domains are indicated. Also, see Figure Supplement 1 and 2.

**Table 1.** Cryo-EM data collection, refinement and validation statistics

|  | **GluD2- in complex with**<br>**7-CKA and Calcium** |
| --- | --- |
| **Data collection and processing** |  |
| Microscope | Titan Krios |
| Voltage (kV) | 300 |
| No. of Micrographs | 2562 |
| Camera | Falcon III |
| Mode of recording | counting |
| Exposure time (seconds) | 60 |
| Electron exposure (e-/Å<sup>2</sup>) | 39 |
| Defocus range (µm) | -1.5 to -3.3 |
| Pixel size (Å) | 1.4 |
| Symmetry | C1 |
| Initial particle images (no.) | 10751 |
| Final particle images (no.) | 7712 |
| Map resolution (Å) | 7.9 |
| FSC threshold | 0.143 |
| **Refinement** |  |
| Initial model used (PDB code) | 5KC8(ATD), 5CC2(LBD) |
| Model resolution (Å) | 7.9 |
| FSC threshold | 0.5 |
| **Model composition** |  |
| Non-hydrogen atoms | 20778 |
| Protein residues | 2616 |
| **R.m.s. deviations** |  |
| Bond lengths (Å) | 0.029 |
| Bond angles (°) | 2.867 |
| **Validation** |  |
| MolProbity score | 1.93 |
| Clashscore | 6.74 |
| **Ramachandran plot** |  |
| Favored (%) | 95.96 |
| Allowed (%) | 3.93 |
| Disallowed (%) | 0.12 |

### The unique architecture of the GluD2 receptor

Our cryo-EM analysis revealed a unique conformation of the tetrameric GluD2 receptor, whereby the Y-shaped arrangement of the extracellular ATD and LBD domains observed in AMPA, Kainate, and GluD1 receptors is disrupted (**Fig. 2 A**). The two dimer arms made up of subunits AB and CD adopt an un-swapped architecture similar to that seen in GluD1 receptors ^15^. The dimer arm AB has two-fold symmetric arrangement at the ATD layer and a quasi-two-fold arrangement of the LBD layer similar to that observed in GluD1 receptors (**Fig. 2 B**). However, in the AB arm, the ATD and LBD dimers are packed on top of each other with their planes, making an angle of ~172°, unlike that in GluD1 where the ATD and LBD planes are placed at an angle of ~99° (**Fig. 2 C**).

**Fig. 2:**
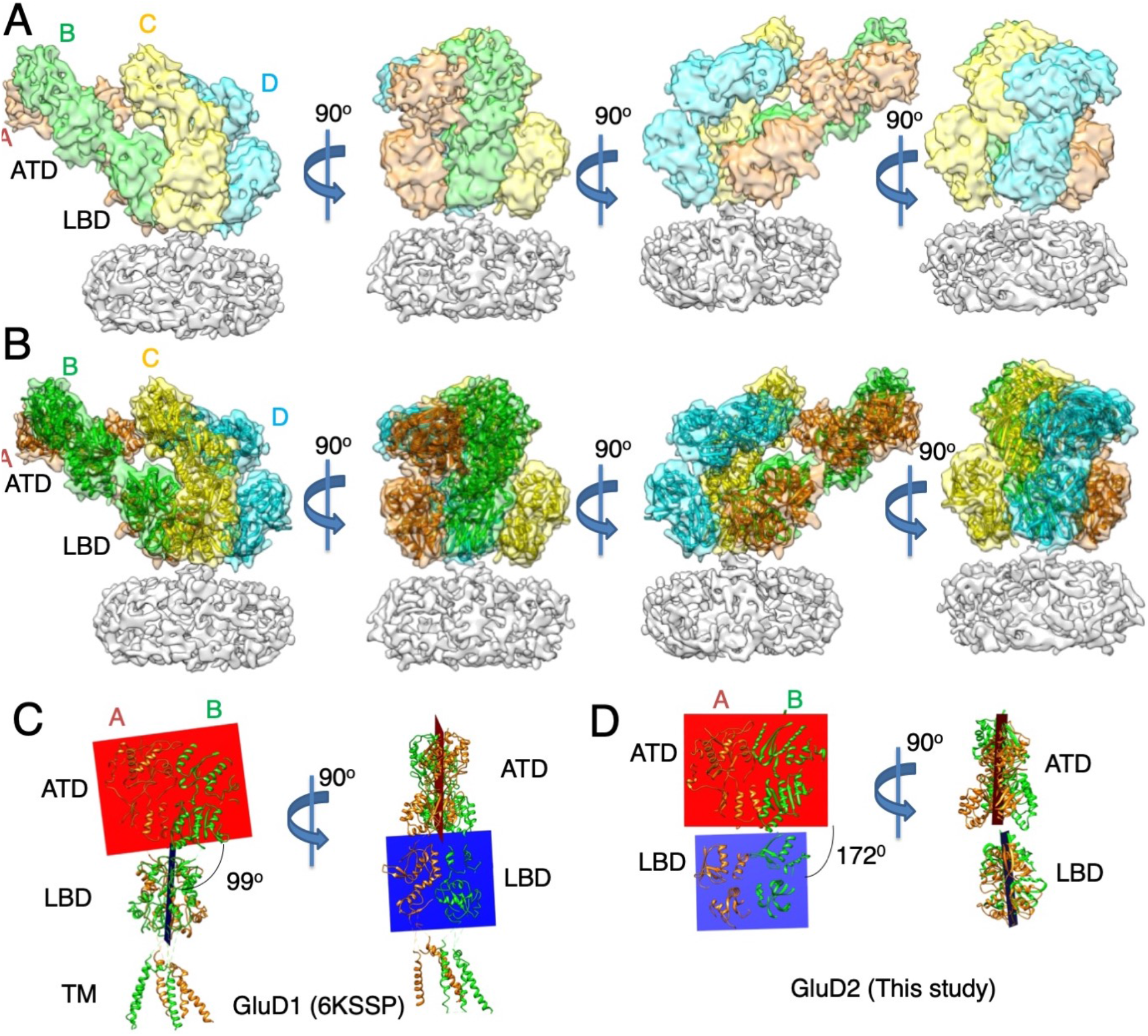
GluD2 has a unique architecture. Panels A and B show the architecture of GluD2 receptor structures determined in the presence of 1mM 7-CKA and 1 mM calcium. **(A)** 3D map of GluD2 where extracellular domains of the receptor tetramer are depicted in different colors corresponding to receptor subunits. TM domains were not resolved and hence are shown in a single grey color. Side view highlighting the broadest face of the Y-shaped receptor and 90° rotated views are shown. The EM map clearly shows the non-swapped arrangement of the ATD and LBD layers, as observed in GluD1. Panel **(B)** shows density map in (A) fitted with protein co-ordinates for the extracellular ATD and LBD domains. Panel C-D show ATD-LBD planes and the angle subtended between them for AB dimer pair of GluD1 (PDB ID: 6KSS) in **(C)** and GluD2 receptors (this study) in **(D).** Also, see Figure Supplement 3.

On the other hand, the CD arm adopts an unusual conformation whereby the ATDs maintain the two-fold symmetry, whereas the LBD dimers are disrupted. The CD arm adopts a bent conformation such that the R1 domains of CD subunits are in close proximity to R2 domains of subunits AB (**Fig. 2 B**). Owing to this, the Centre of mass (COMs) of R1 and R2 lobes of subunit-A subtend an angle of ~136° with COM of R1 lobe of ATD from subunit C. Similarly, the R1 and R2 COMs of subunit-B makes an angle of ~111° with R1-COM of ATD from subunit D. Taken together, this arrangement of ATD and LBD domains results into a unique asymmetric architecture of the extracellular domains of GluD2 receptor (**Supplementary Fig. 3**).

### Organization of the Amino-terminal domains

Consistent with the nano-molar affinity of the GluD2 ATDs for dimerization ^12^, they exist in a 2-fold symmetric dimeric configuration (**Fig. 3 A**). However, owing to the unique asymmetric arrangement of the two extracellular dimer arms of GluD2, they lie in different planes. The distances between the Center of Mass (COMs) of the ATDs between subunits AB and CD are ~34 Å and ~35 Å consistent with the dimeric arrangement. However, the distance between COMs of subunits A and D and B and C are ~76 Å and ~64 Å due to asymmetric placement of the CD dimer pair (**Fig. 3 B**). Moreover, we also observe that in GluD2 ATD dimer packing is not compact (buried surface area of ~1647Å^2^ for AB dimer pair) like that observed in GluD1 (buried surface area of ~2120Å^2^) as evidenced by the buried surface area analysis. Due to this asymmetric placement of ATDs, the COMs of R2 lobes of ATD subunits A-D are at a distance of ~73 Å, and an angle of ~59° is formed between COMs of subunits A, D, and C. Further, COMs of R2 lobes of ATD subunits B and C are at a distance of ~63Å and make an angle of ~125° with COM of subunit D (B-C-D) highlighting the unique asymmetric arrangement of the ATD layer (**Fig. 3 C**).

**Fig. 3:**
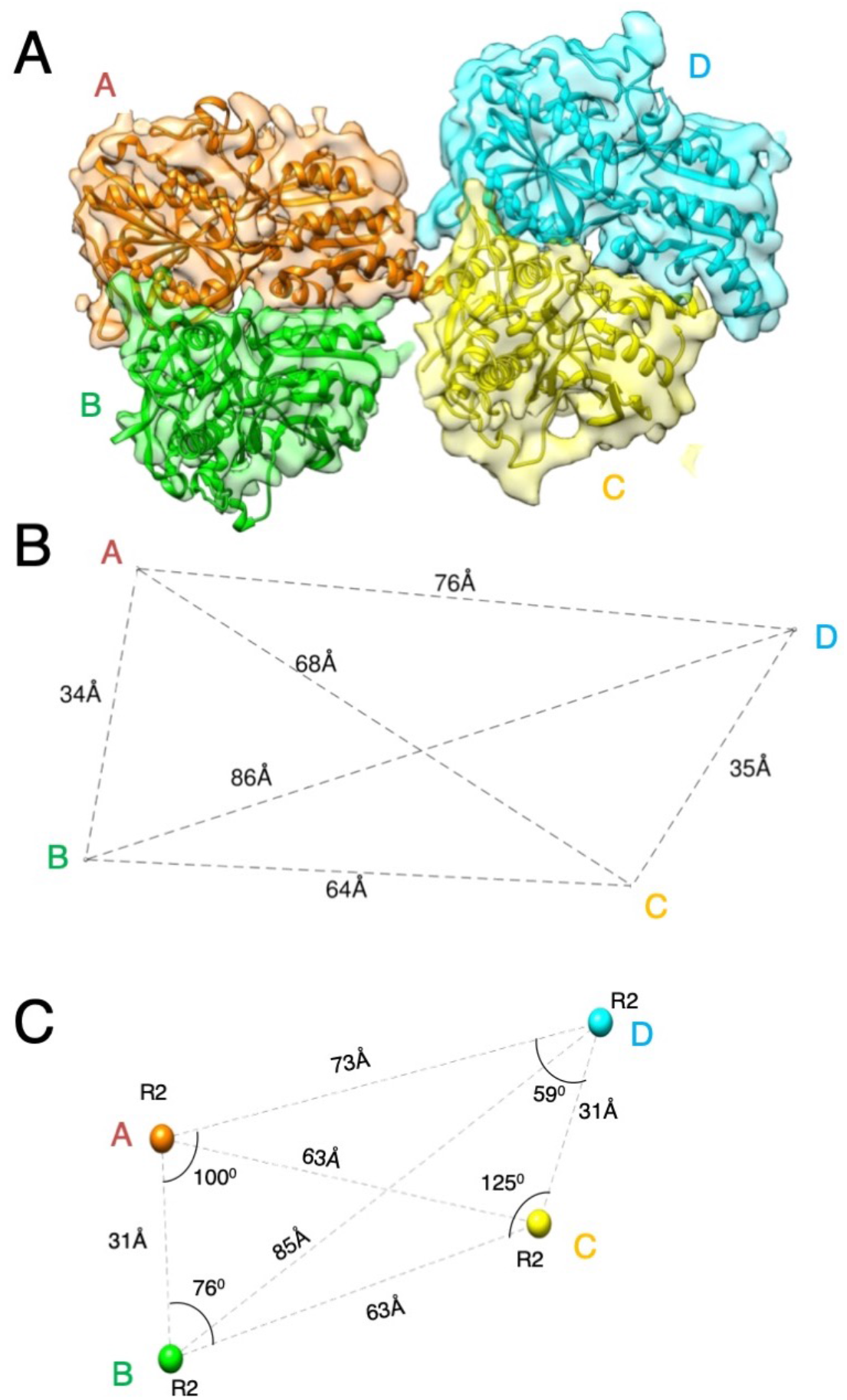
Arrangement at the ATD layer. **(A)** The top view of the ATD domains fitted in the segmented density map is shown. The subunit colors are depicted as in Figure 2. **(B)** The distances between the Center of Mass (COMs) of the ATD domains are shown. Panel (C) shows distance and angles subtended between COMs of R2 lobes of the ATD domains.

### Orientation and arrangement of the LBD domains

Unlike ATDs where the dimeric arrangement is conserved for both the extracellular arms of the receptor tetramer, the arrangement at the LBD layer is unique. While subunits A and B maintain a quasi-two-fold dimeric arrangement, the subunits C and D adopt a desensitized-like state with LBD dimers disrupted (**Fig. 4 A**). Consistent with this LBD arrangement, the distances between the Center of Mass (COMs) of the LBDs between subunits A-B and C-D are ~34 Å and ~50 Å. However, the distance between subunits A and D and B and C are ~60 Å and ~42 Å due to asymmetric placement of the C and D subunit LBDs (**Fig. 4 B**). Moreover, the buried interface for the AB-LBD dimer pair is ~341 Å^2^ in contrast to ~1184 Å^2^ observed in the corresponding LBD dimer of GluD1 further highlighting the weak dimer packing. However, the distances between the COMs of D1 and D2 domains of LBDs are ~25 Å and ~26 Å for subunits A-B and C-D respectively (**Fig. 4 C**). This is similar to that observed in the GluD1 receptor structure determined in the presence of same ligands (1mM Ca^2+^ and 7-CKA) ^15^ indicating similar cleft closure.

**Fig. 4:**
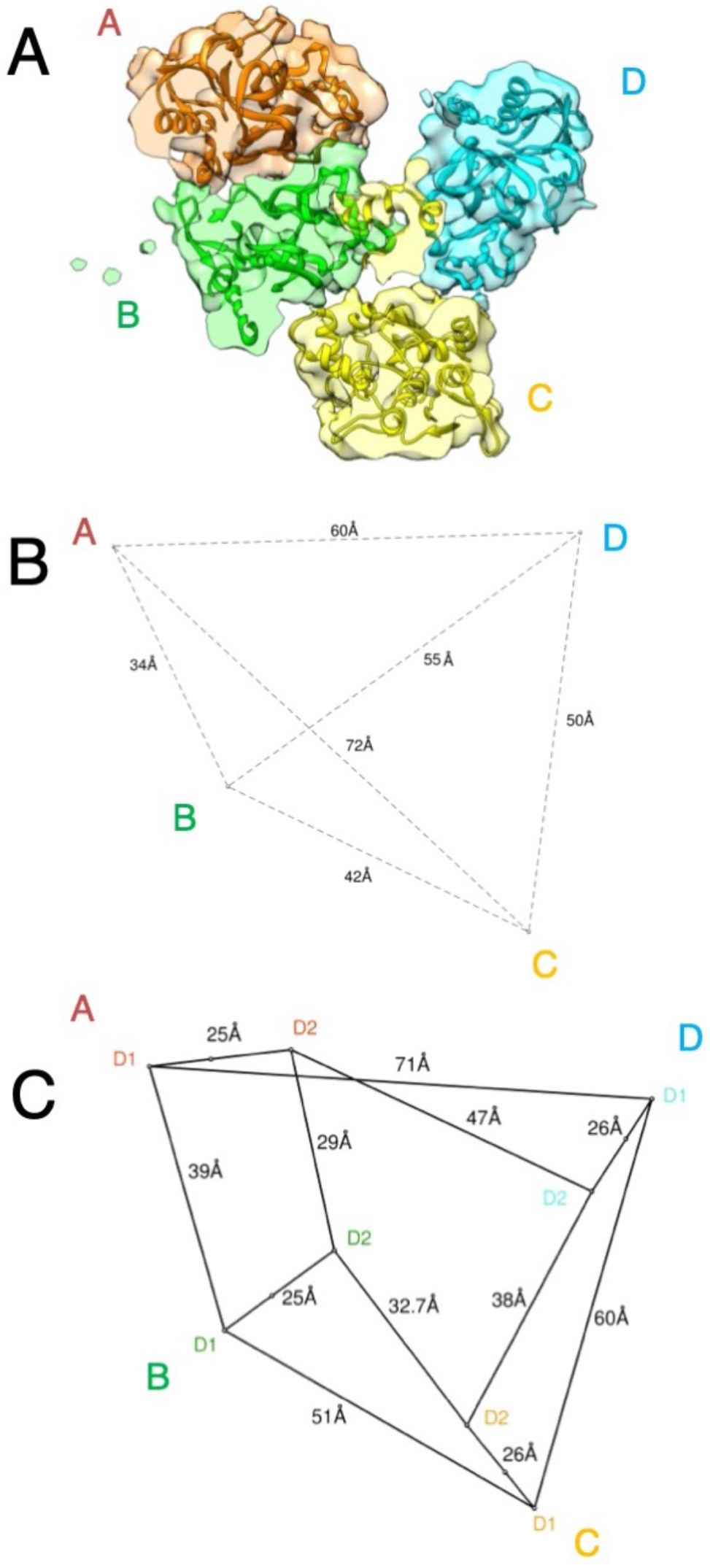
Orientation and arrangement of the LBD domains. **(A)** The top view of the GluD2 LBD domain is shown fitted into the segmented EM map. While subunits A and B maintain a quasi-two -fold dimeric arrangement, the subunits C and D adopt a desensitized -like state with LBD dimers disrupted. **(B)** The distances between COMs of the LBD domains are shown by dashed lines. **(C)** shows the distance between COMs of D1 and D2 lobes of the LBD domains for AB dimer and CD subunits. Also, see Figure Supplement 4.

## Discussion

We utilized a GluD2 construct with minimal modification (only C-terminal truncation) for expression, purification and structure determination by cryo-EM. Although the resolution of our structure is ~7.9 Å and the transmembrane domains are not resolved, we were able to generate model for the extracellular domains of the receptor by fitting crystal structures of ATD and LBD domains into the constraints of EM density map. The GluD2 structure reveals a non-swapped arrangement of the ATD and LBD domains like GluD1 receptors and reconfirms that this tetrameric receptor arrangement is unique to ‘orphan’ delta glutamate receptor family. The structural differences of GluD2 with GluD1 in similar imaging conditions could point towards unique functional properties of GluD2. However, we cannot rule out the influence of grid preparation conditions and air-water interface interactions in observation of this unique asymmetric arrangement of the two-extracellular arms. Moreover in the absence of domain swapping, the conformational freedom for the movement of the two dimer arms is substantially higher as seen in the GluD1 receptors ^15^ and unlike that in AMPA ^20 21^ and kainate receptors ^22 23^ and we could have trapped a physiologically relevant conformational intermediate. It’s also important to note that this conformation may not be compatible with trans-synaptic, tripartite complex formed between GluD2, cerebelin, and neurexin which is essential for maintaining the synaptic integrity of PF-PC synapses as this would limit the large-scale motions of the two extracellular arms. Thus it’s likely that in absence of the stabilizing trans-synaptic interactions and the conformational mobility due to non-swapped architecture, receptor may adopt the conformation seen in our structure.

The non-swapped architecture observed in the orphan delta receptor family, short ATD-LBD linkers, and weak ligand affinity might be some of the reasons behind receptor inactivity. Furthermore, the non-swapped architecture might be dictated by the shorter linker between the ATD-LBD. However, more experiments are required to establish this notion. In overview, GluD2 structure presents a framework that could pave way for understanding the enigmatic properties of these orphan receptors.

## Methods

### Construct design

Rat GluD2 was cloned into the pEGBacMam vector ^24^ in frame with a C-terminal thrombin recognition site (GLVPRGSAAAA) and EGFP (A207K mutant) with a C-terminal octahistidine (His8) tag. Full-length GluD2 had weak expression and stability as judged via FSEC ^16^. Further screening of constructs identified GluD2Δ870 as a promising candidate for overexpression and purification. All the constructs were verified by sequencing of the entire coding region.

### Expression and Purification of GluD2

The HEK293S GnTI^-^ (ATCC CRL-3022) were obtained and authenticated by ATCC, and no further authentication or mycoplasma testing was performed. Expression and purification were carried out as reported earlier ^15^. Briefly, suspension adapted HEK293GnTI^-^ cells growing in freestyle 293 expression media supplemented with 2 % FBS (Gibco), 2 mM Glutamine (Gibco) and 1% Penstrep (Gibco) and were infected with high-titer baculovirus having the GluD2 construct at a multiplicity of infection (MOI) of ~1. To boost the expression of the protein, 10mM sodium butyrate (Sigma) was added 20 hrs post-infection and cultures moved to 30°C. The cells were harvested ~48-52 hrs later, disrupted, and membranes were fractionated. The membrane was detergent-solubilized in a buffer containing 150 mM NaCl, 20 mM Tris (pH 8.0), 40 mM n-dodecyl-β-D-maltopyranoside, and 6 mM cholesterol hemisuccinate at 4°C. IMAC purification using cobalt-charged TALON metal affinity resin was carried out. Protein fractions were pooled, and thrombin digested overnight to cleave off the EGFP and His tag. The thrombin digested protein was further purified by size exclusion chromatography (Superose 6 10/300) equilibrated with 150 mM NaCl, 20 mM Tris pH 8.0, and 0.75 mM DDM, 0.03 mM cholesterol hemisuccinate. Eluted fractions were analyzed for homogeneity by SDS-PAGE and FSEC, only fractions containing tetrameric receptors were pooled and concentrated to ~0.8 mg/ml.

### Cryo-electron microscopy data collection and analysis

A droplet of 3 μl of purified GluD2 Δ870 at a concentration of ~0.8 mg/ml was applied onto a glow discharged 1.2/1.3 300 mesh ultra foil gold grid (Quantifoil). It was blotted for 3 seconds at a blot force of 0 using the FEI Vitrobot Mark IV at 14°C and 95 % humidity, and plunge-freezing the grid in liquid ethane. Cryo-EM data collection was carried on a 300 kV Titan Krios microscope equipped with a Falcon 3 camera. Micrographs were recorded in super-resolution mode at a magnified nominal pixel size of 1.4 Å and defocus ranging from −1.5 to −3.3 μm. Each micrograph consisted of 25 dose-fractionated frames with a total exposure time of 60 s and a total dose of 39.0 e-per Å^2^. A total of 2562 movies were collected, aligned, and dose-weighted to correct for movement during imaging and account for radiation damage via Motioncor2 ^25^. The CTF parameters for each micrograph were determined by Gctf ^26^. ~61000 particles were picked manually and subjected to several rounds of reference-free 2D classification followed by manual inspection and selection of classes with iGluR like features. This process yielded a stack of ~10751 cleaned particles that were subjected *ab initio 3D* reconstruction in cryoSPARC V2 ^18^ into two classes. Class two with 7712 particles was subjected to further refinement in cryoSPARC v2.12.4 using the new features of aberration corrections and per-particle defocus refinement as implemented in the new homogenous refinement module. Homogenous 3D refinement in C1 symmetry, followed with the non-uniform ^27^ and local refinement, yielded a final map to a resolution of ~ 7.9 Å (0.143 FSC). The final 3D map was sharpened and used for model building. Masked refinement (ATD-LBD) gave better resolution as per FSC at 0.143, however, the map quality didn’t improve.

### Model building and refinement

GluD2 tetramer model was built by the rigid-body fitting of individual ATD and LBD domains into the EM map in UCSF Chimera ^28^. Four copies each of GluD2 amino-terminal domain (PDB code, 5KC8) and ligand-binding domain (PDB code, 5CC2) was used. Owing to weaker EM density, the TM domain was not modeled. The fits were improved by using a molecular dynamics-based flexible fitting simulation ^29^ followed by real-space refinement in Phenix ^30^. Model-map FSC curve calculation yielded values of ~7.9 that agreed well with the gold-standard FSCs generated during the 3D refinement (**Figure Supplement 1**). The final model has good stereochemistry, as evaluated using MolProbity ^31^ (**Table 1**). All of the figures were prepared with Pymol ^32^ and UCSF Chimera ^28^.

### Statistics

No statistical methods were used to predetermine the sample size. The experiments were not randomized, and the investigators were not blinded to allocation during experiments and outcome assessment.

### Data Availability

The cryo-EM density reconstruction and final models were deposited in the Electron Microscopy DataBase (accession codes EMD-XXXX) and the Protein Data Bank (accession codes XXXX). All other relevant data supporting the key findings of this study are available within the article and its <u>Supplementary Information</u> files or from the corresponding author upon request.

## Acknowledgments

This work was supported by the Wellcome Trust DBT India Alliance fellowship (grant number IA/I/13/2/501023) awarded to J. Kumar. A. P. Burada thanks ICMR, India, for senior research fellowship. Dr. M. L. Mayer, NIH, Bethesda kindly gifted the various iGluR constructs that were sub-cloned and used for construct optimization and mutational studies. Dr. E. Gouaux, OHSU, Portland kindly provided the pEG BacMam vector. Access to EM was provided by the National Electron cryo-microscopy facility at the Bangalore Life Sciences Cluster. Funding for this was provided by a grant from the Department of Biotechnology, Government of India, DBT/PR12422/MED/31/287/2014. We thankfully acknowledge the kind assistance of Dr. Vinothkumar Kutti Ragunath, NCBS, Bangalore in grid preparation, and EM data collection.

## Author Contributions

A.P.B optimized construct, expressed, purified protein, and processed EM data. JK supervised the overall project design and its execution. Both the authors contributed to the analysis and preparation of the manuscript and approved the final draft.

## Corresponding author

Correspondence to Janesh Kumar

---

**Supplementary Fig. 1:**
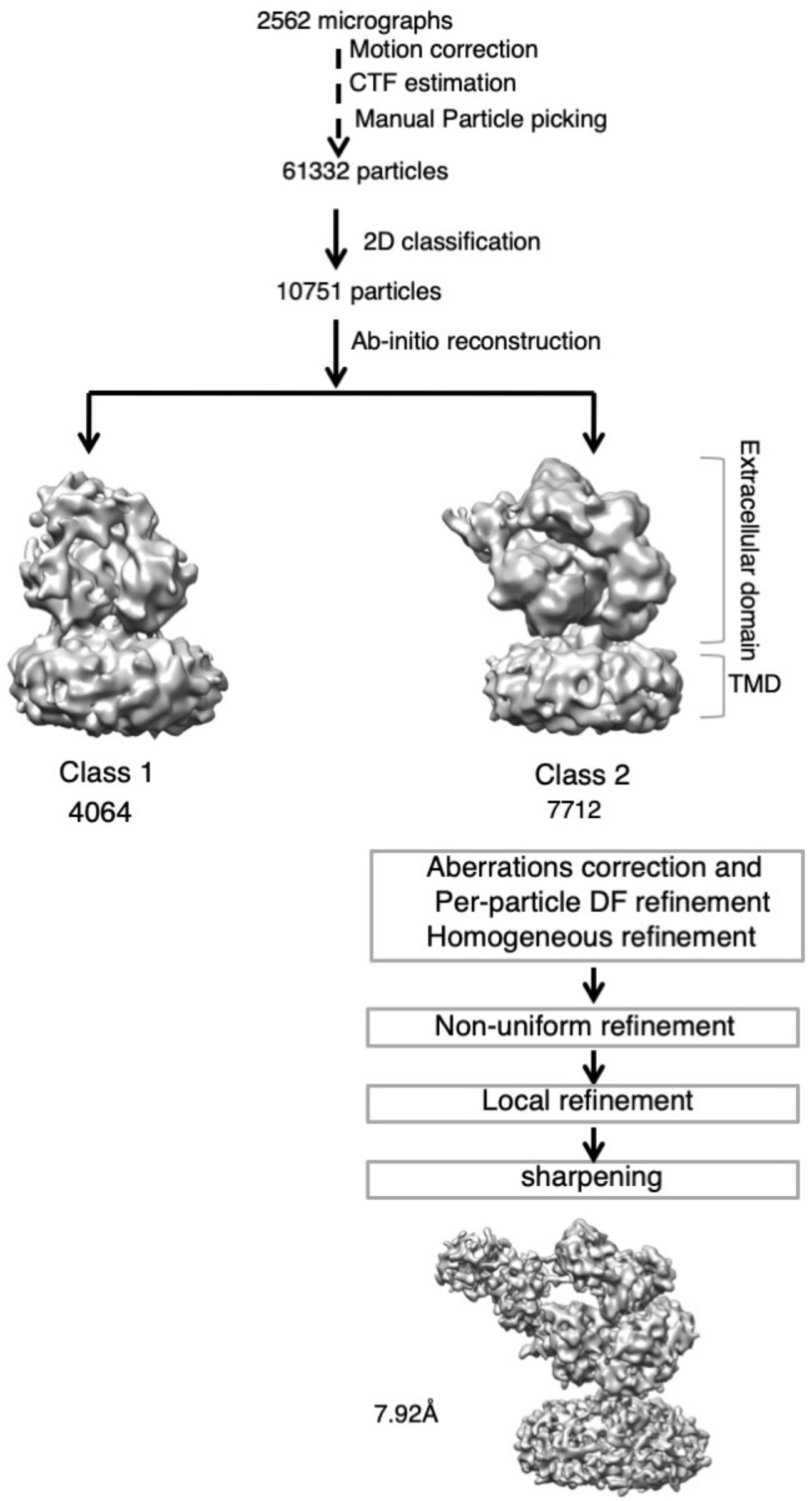
Cryo-EM data processing workflow. A schematic of Data processing workflow is shown.

**Supplementary Fig. 2:**
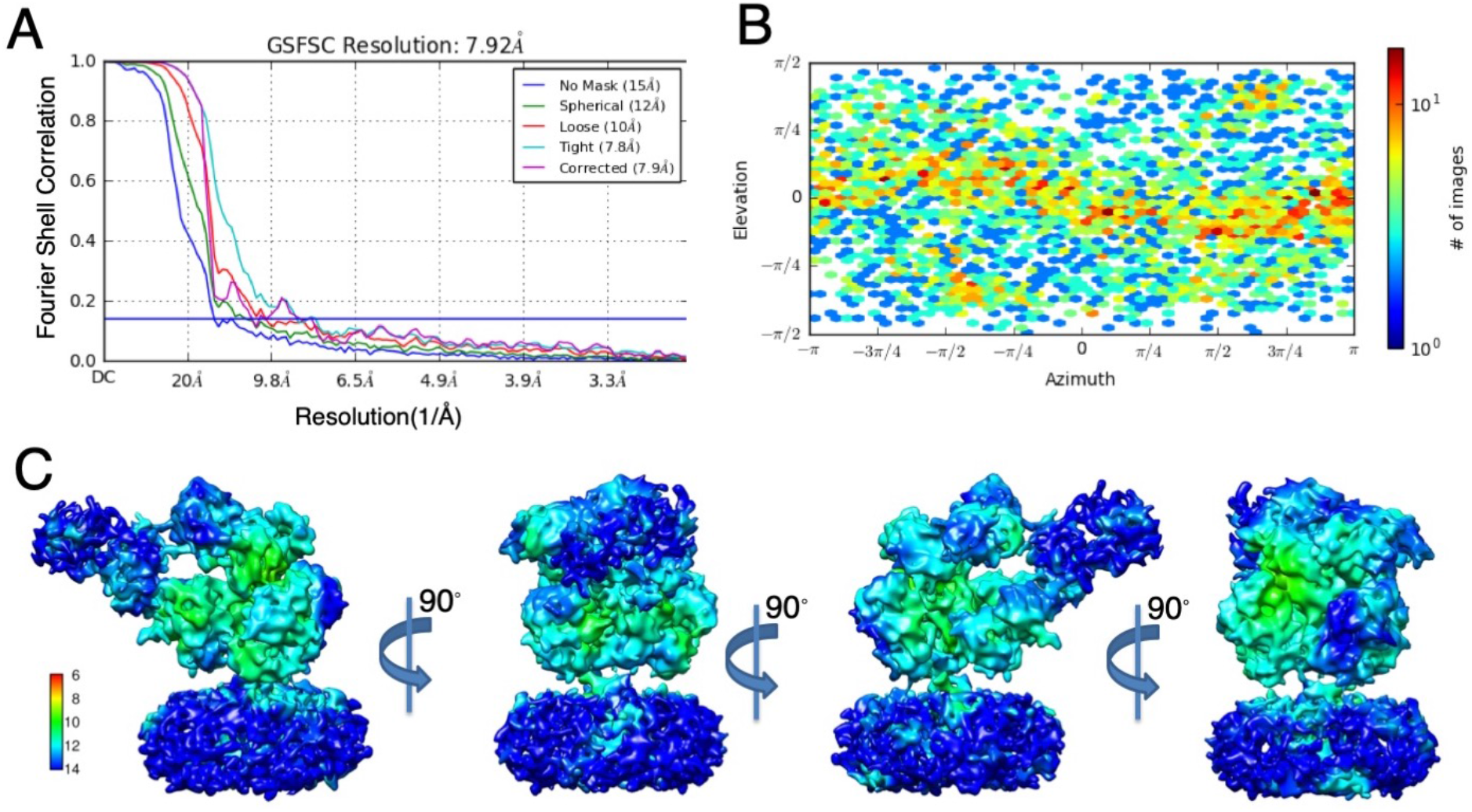
Local resolution estimates of the GluD2 cryo-EM map. **(A)** Fourier shell correlation curves for the final 3D refinement run. The resolution of the map corresponding to FSC 0.143 is indicated. **(B)** Euler angle distribution of particles for the 3D model is shown. **(C)** The sharpened Cryo-EM map of GluD2-870 in 7-CKA and calcium bound form colored based on local resolution.

**Supplementary Fig. 3:**
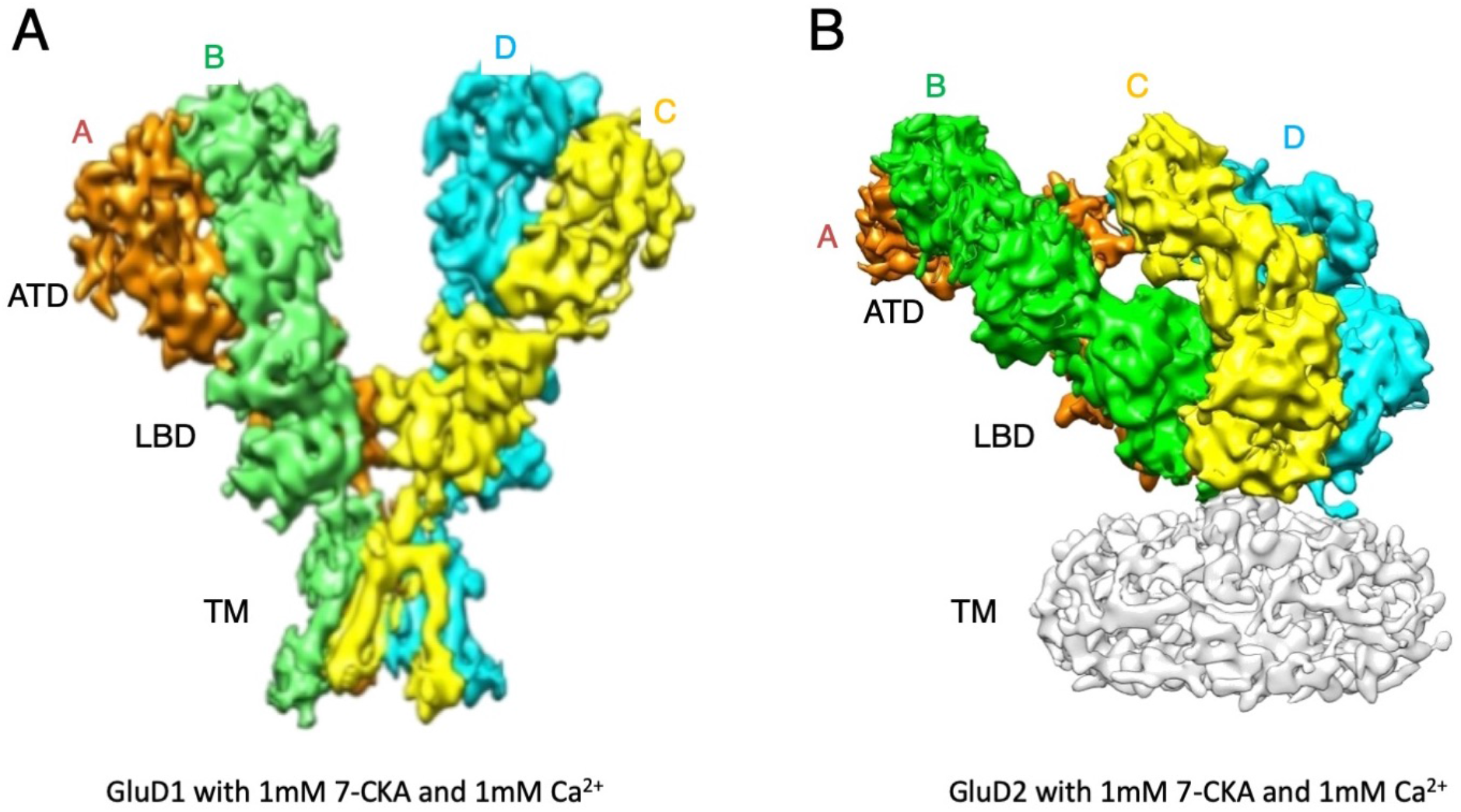
GluD1 and GluD2 structures in presence of calcium ions and 7-CKA. GluD1 (PDB ID: 7KSP) **(A)** and GluD2 **(B)** are shown with the broadest view parallel to the membrane highlighting the arrangement of the extracellular domains.

